# eDNA in a bottleneck: obstacles to fish metabarcoding studies in megadiverse freshwater systems

**DOI:** 10.1101/2021.01.05.425493

**Authors:** Jake M. Jackman, Chiara Benvenuto, Ilaria Coscia, Cintia Oliveira Carvalho, Jonathan S. Ready, Jean P. Boubli, William E. Magnusson, Allan D. McDevitt, Naiara Guimarães Sales

## Abstract

The current capacity of environmental DNA (eDNA) to provide accurate insights into the biodiversity of megadiverse regions (e.g., the Neotropics) requires further evaluation to ensure its reliability for long-term monitoring. In this study, we first evaluated the taxonomic resolution capabilities of a short fragment from the 12S rRNA gene widely used in fish eDNA metabarcoding studies, and then compared eDNA metabarcoding data from water samples with traditional sampling using nets. For the taxonomic discriminatory power analysis, we used a specifically curated reference dataset consisting of 373 sequences from 264 neotropical fish species (including 47 newly generated sequences) to perform a genetic distance-based analysis of the amplicons targeted by the MiFish primer set. We obtained an optimum delimitation threshold value of 0.5% due to lowest cumulative errors. The barcoding gap analysis revealed only a 50.38% success rate in species recovery (133/264), highlighting a poor taxonomic resolution from the targeted amplicon. To evaluate the empirical performance of this amplicon for biomonitoring, we assessed fish biodiversity using eDNA metabarcoding from water samples collected from the Amazon (Adolpho Ducke Forest Reserve and two additional locations outside the Reserve). From a total of 84 identified Molecular Operational Taxonomic Units (MOTUs), only four could be assigned to species level using a fixed threshold. Measures of α-diversity analyses within the Reserve showed similar patterns in each site between the number of MOTUs (eDNA dataset) and species (netting data) found. However, β-diversity revealed contrasting patterns between the methods. We therefore suggest that a new approach is needed, underpinned by sound taxonomic knowledge, and a more thorough evaluation of better molecular identification procedures such as multi-marker metabarcoding approaches and tailor-made (i.e., order-specific) taxonomic delimitation thresholds.

## INTRODUCTION

The need for advancing our understanding of the world’s biodiversity increases in parallel with the acceleration of anthropogenic impacts on the planet’s ecosystems. To implement strategies to minimise the effects of human impacts, understanding compositions of species assemblages within ecosystems is paramount (Morris, 2010). This task is particularly difficult when investigating megadiverse regions of the world such as the Neotropics, which harbour an extremely large diversity of living organisms. The Amazon basin, for example, is estimated to hold the highest diversity of freshwater fish found anywhere on the planet (Albert & Reis, 2011). To date, it has been documented that 2,406 species belonging to 514 genera and 56 families of fish inhabit the tributaries of the Amazon River, with many more not yet described (Jézequel et al., 2020). Undoubtedly, this region serves as a biodiversity hotspot, with Amazonian fishes representing ~15% of all freshwater fish species described worldwide (Jézequel et al., 2020; Leroy et al., 2019).

Due to the increase in anthropogenic impacts in Neotropical rivers (e.g., pollution, siltation, mining and damming), there is a growing danger that this rich biodiversity will be lost before it can be fully described (Alho, Reis & Aquino, 2015; Agostinho, Thomaz & Gomes, 2005). This emphasizes the urgency of accurate and rapid biodiversity assessments throughout the region. Although years of biomonitoring in the Neotropical region have been conducted, inventories of fish fauna remain incomplete (Frota, Depra, Petenucci & Graça, 2016) demonstrating the need for improvements in biodiversity assessment methods through a more integrative approach aimed at circumnavigating traditional sampling limitations.

A powerful addition to biodiversity surveying is the application of DNA-based approaches. The use of environmental DNA (eDNA; i.e., DNA extracted from environmental samples such as water or soil; Taberlet et al., 2012) for biomonitoring is now widespread, particularly within freshwater ecosystems (Hering et al., 2018; Senapati et al., 2018). Advances in next-generation sequencing (NGS) have unlocked the potential use of eDNA metabarcoding for monitoring whole communities within specific taxonomic groups (e.g., fishes; Miya, Gotoh & Sado, 2020). Recent studies display its efficiency in different aquatic environments and show how it compares favourably to, or even outperforms, traditional sampling methods in terms of species detections (McDevitt et al., 2019; McElroy et al., 2020) and facilitates investigations into patterns of extirpation, invasive species detection, and dynamics of species richness (Lacoursière-Roussel et al., 2018; Sales et al., 2021). Despite the increase of eDNA surveys in megadiverse systems, several limiting factors prevent its full application (Cilleros et al., 2019; Doble el al., 2020; Sales et al., 2021). Although challenges associated with the collection and preservation of samples have already been addressed (Sales et al., 2019), obstacles in taxonomic assignments remain understudied.

The ability to detect species is reliant on the reference library used to assign retrieved sequences to species level and on the robustness of the taxonomic resolution of targeted gene fragments (Sassoubre, Yamahara, Gardner, Block & Boehm, 2016). As outlined by the International Barcode of Life initiative (Hebert, Cywinska, Ball & Dewaard, 2003), databases targeting mitochondrial cytochrome oxidase subunit I (COI) are generally more complete than other gene regions such as the mitochondrial 12S and 16S rRNA genes. However, the COI region has been shown to be less suitable for eDNA metabarcoding work for vertebrates (due to inflated detections of non-target organisms), therefore substantiating the need to explore the use of more suitable gene regions such as 12S and 16S and to expand their respective databases (Collins et al., 2019; Deagle, Jarman, Coissac, Pompanon & Taberlet, 2014).

In this context, a set of universal PCR primers for metabarcoding eDNA from fishes has been developed targeting a length of around 163-185 bp of the 12S rRNA gene region (MiFish; Miya et al., 2015). This widely used primer set has been pivotal in describing fish communities on a truly global scale (Miya, Gotoh & Sado, 2020), and has also provided meaningful information for species-rich rivers (Ahn et al., 2020). Despite the efforts made by global barcoding initiatives towards the development of more comprehensive reference databases, in most circumstances these databases remain far from complete, especially for the currently commonly used 12S mitochondrial gene region (Doble et al., 2020; Sales et al., 2020; Weigand et al., 2019). Furthermore, the proportion of species sequenced is lower in species-rich regions, such as the Neotropics with poorly sampled habitats and taxa. As it stands now, only a limited number of fish species can be found in DNA databases, hindering the potential of eDNA metabarcoding as a biomonitoring method in these regions (Cilleros et al., 2019; Sales et al., 2019, 2021).

Besides the well-known obstacles posed by incomplete reference databases, the taxonomic resolution of target amplicons can hinder inferences of species occurrence. For example, the current fixed general threshold used for species assignment (e.g., > 97% similarity - Sales et al., 2021; >98% similarity - Marques et al., 2020) assumes the existence of a barcoding gap (i.e., presence of a gap between the highest intraspecific and the lowest interspecific variation within the analysed taxonomic group; Meyer & Paulay, 2005). The accuracy of selected markers in detecting species relies on the separation between the intra and inter-specific divergences, and the greater the overlap between these variations, the less effective DNA barcoding becomes (Meyer & Paulay, 2005). In the absence of a barcoding gap, the use of a general threshold may lead to an inaccurate taxonomic assignment, over splitting some taxa while lumping together others. This issue is particularly important in highly biodiverse areas, especially where the proportion of recently diverged species and cryptic species is relatively high. In this regard, Sales et al. (2021) have highlighted issues of low taxonomic resolution with the widely used 12S MiFish marker, unable to assign members of the genus *Prochilodus* to the species level. This matter then raises conservation issues as native species (e.g., *P. hartii*) could be wrongfully assigned to congeneric invasive species (e.g., *P. argenteus*). eDNA metabarcoding represents a technological leap for characterizing and assessing biodiversity (Petruniak, Bradley, Kelly & Hanner, 2020), but these obstacles can represent a bottleneck to its application in highly biodiverse regions (Sales et al., 2020, 2021).

In this study, we aim to assess the use of eDNA metabarcoding as a tool to estimate fish biodiversity in a megadiverse Neotropical system, by directly comparing it to data obtained by netting surveys carried out at the same time and sites. We adopted a step by step, integrative approach using newly generated and existing sequence data from a wide range of neotropical fish species, eDNA from water samples and netting data. We first performed a genetic distance-based analysis to investigate the optimum delimitation values based on the 12S MiFish primers, followed by an assessment of the performance of the delimitation values through a barcoding gap analysis. We then analysed water samples collected from the Brazilian Amazon and we compared measures of α-diversity (species richness) and β-diversity (change in species composition among locations) generated from eDNA and traditional netting data collected in the same sampling sites.

## METHODS AND MATERIALS

### Evaluation of taxonomic resolution power of the 12S rRNA fragment

In order to improve taxonomic assignment, 47 fish species (Table S1) were sequenced and included in a customised reference database for Neotropical fishes. Tissue samples were provided by the Grupo de Investigação Biológica Integrada (GIBI) tissue collection, located at the Universidade Federal do Pará (UFPA; Belém, Brazil). Fragments of the mitochondrial 12S rRNA gene were obtained using PCR (conducted using MiFish primers, following the same protocols used for eDNA samples described below) and Sanger sequenced. Consensus sequences were obtained with Geneious v8.1 (Kearse et al., 2012).

Several analyses were conducted to verify the taxonomic resolution of 12S rRNA fragments targeted by the MiFish metabarcoding primers. For this stage of the analysis we created a combined enhanced dataset comprising 264 neotropical fish species, belonging to eight orders (Table S2) from existing GenBank and newly sequenced data (373 barcodes in total). Threshold optimization analysis was conducted using the SPIDER package in R v3.5.1 (R Core Team 2019) through the *threshOpt* function. This returns the false positive and negative rates of identification accuracy for different threshold values as well as providing the total cumulative errors (false positive + false negative). When applying a range of thresholds, this function allows the optimization of values aiming to minimise the error rates. The default threshold for this function is set to 0.01 (1%), however, this can be changed using the *threshVal* function, and here were included values ranging from 0.001 to 0.03 (0.1% to 3%). Using the thresholds generated from the *threshVal* function, a genetic-based delimitation analysis was performed using K2P genetic distances (Kimura, 1980). The threshold estimates were applied as the best delimitation values to estimate species. Milan et al. (2020) suggested threshold values ranging from 0.4% to 0.55% for a different fragment of the 12S rRNA gene. Currently, no delimitating reference values for the MiFish 12S marker used here have been published, thus we established distance thresholds, which were then used within the *threshID* function. The *threshID* function assigns four possible results for each sequence in the dataset: “correct”, “incorrect”, “ambiguous”, and “no ID”. The “correct” results suggest that all matches within the threshold value of the query are the same species and “no ID” shows that no matches were found to any individual within the threshold.

SPIDER was also used as a means to investigate the presence/absence of the “barcoding gap” by identifying the furthest intraspecific distance among the same species, using the *maxInDist()* function and the closest non-conspecific using the *nonConDist()* function. The occurrence of no barcoding gap is represented by a zero or negative distance as a result of the maximum intraspecific distance being subtracted from the minimum interspecific distance.

### Study sites and sample collection

The eDNA component of the study was conducted in six different sites located in the Brazilian Amazon, with four sites inserted inside the Adolpho Ducke Forest Reserve (sites B1-B4; Fig. 1; Table S3) and two outside (A and C). The Reserve is a designated 100 km^2^ area protecting continuous and non-isolated rainforest established by the National Institute of Amazon Research (INPA, for a more detailed description see Supp. Information). The Ducke Reserve represents one of the first sites of the Brazilian Long-Term Ecological Research Program (PELD) and the Biodiversity Research Program (PPBio), run by the Brazilian Ministry of Science and Technology. A long-term study by Zuanon et al. (2015) has revealed that the streams of the Ducke Reserve comprise an estimated 70 species of fish from seven taxonomic orders: Characiformes, Siluriformes, Gymnotiformes, Perciformes (from which now Cichliformes have been separated), Cyprinodontiformes, and Synbranchiformes. The most diverse taxonomic order in the reserve is the Characiformes, comprising six families and 24 species followed by the Siluriformes with seven families and 17 species, Gymnotiformes with four families and 12 species, Perciformes with two families and 13 species, Cyprinodontiformes with one family and three species, and Synbranchiformes with one family and one species.

**Figure 1.**
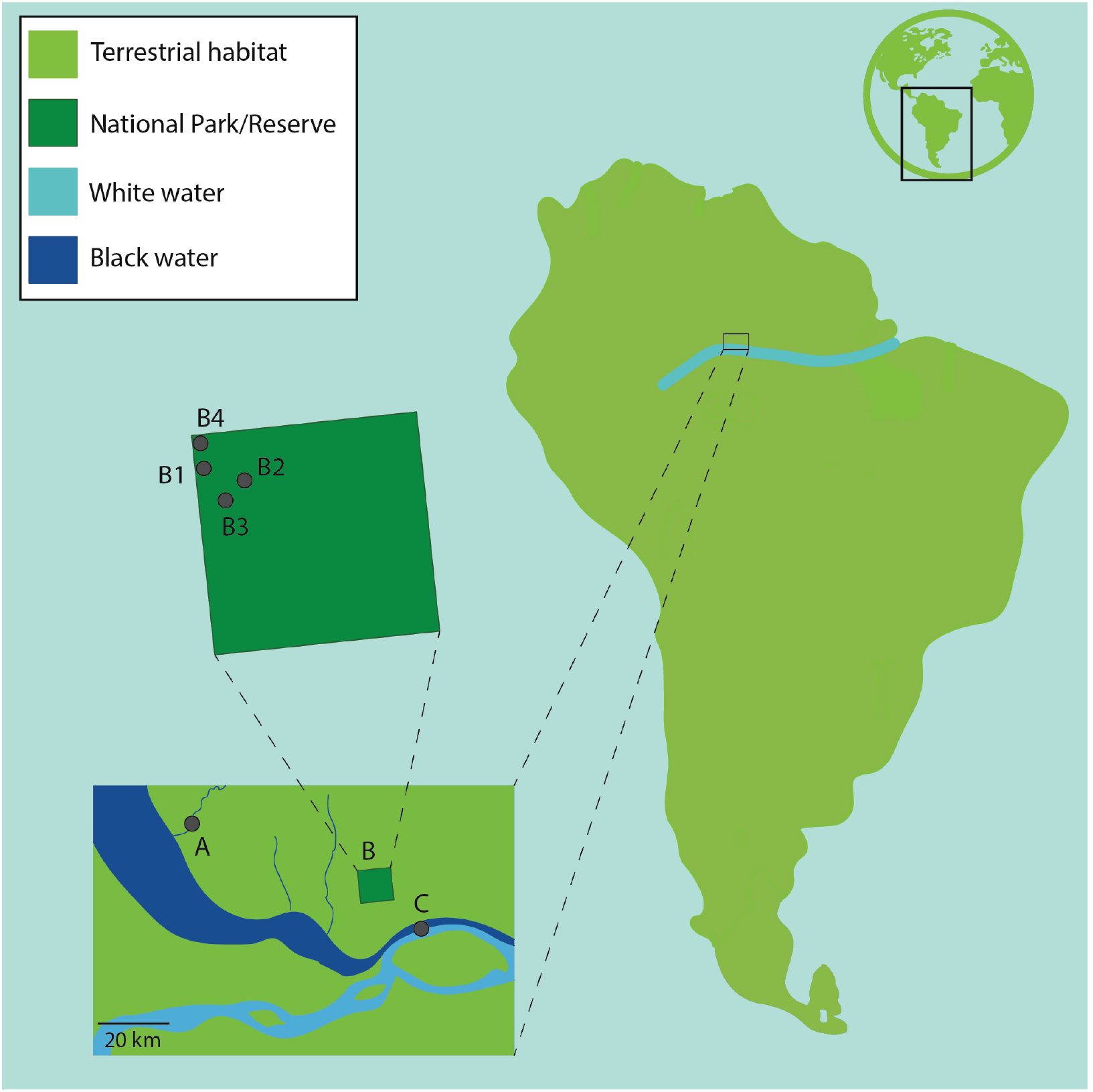
Sampling sites located in the Adolpho Ducke Forest Reserve (B1-B4), and the two additional sites located outside the Reserve (A: Aturiá stream, and C: confluence between the Solimões and Negro rivers - *“the meeting of the waters”*).

Two main drainage basins have there headwaters near the centre of Ducke Reserve, one on the western side of the reserve that flows to the black waters of the Rio Negro and one on the eastern side that flows to the white (sediment laden) waters of the Amazon River. Sampling within the reserve was carried out on the north-west side, along four tributaries of Acará. Three sites were on unnamed third-order streams (B1-B3; Fig. 1) and one was on Barro Branco, a second-order stream (B4; Fig 1). Net sampling and eDNA sampling protocols were both carried out in January 2019. Additionally, for the eDNA analysis, two more sites were sampled opportunistically outside the Reserve (A and C; Fig. 1), one on Aturiá stream, which is a northern tributary of the Rio Negro (A; Fig. 1), and the other at “the meeting of the waters” where the black waters from the Rio Negro and the white waters of the Solimões (upper Amazon) form the Amazon River (C; Fig. 1).

For the eDNA metabarcoding, from each of the four streams within Ducke Reserve, three water replicates were taken from three positions (from the left bank, middle and right bank) along a transect at 0m, 25m, and 50m using 500 ml water bottles resulting in a total of nine replicates per stream. Two water-sample replicates were collected at site A and three at site C (no netting was performed at these sites). At the start of each sampling period, a field blank was collected, totaling four field blanks overall. Samples were collected prior to the start of the netting sampling to avoid the risk of contamination. Water samples were manually filtered using a syringe and Sterivex enclosed filters (0.45μm, Merck Millipore) and kept cool until transport to the UK. In total, 51 samples were analysed (including 41 water samples, four field blanks, two extraction blanks, four PCR negative controls, Table S4). The eDNA extraction, amplification of the 12S rRNA fragment using the MiFish primer set, and library preparation were conducted following the procedure described in Sales et al. (2021).

Netting was conducted in the same four sites in the Reserve (sites B1-B4) straight after water was collected for eDNA analysis (McDevitt et al., 2019), spanning a range of 50m. Each stream was sampled following the rapid assessment protocol RAPELD (Magnusson et al., 2005). Two nets (5 mm mesh size) were placed at the edge of the sampling area to prevent fish accessing and exiting the study area. An additional net was used to subdivide the full lenght in segments. A total of three sub-stretches of ~16 meters each moving upstream were covered. The fishes were collected with nets and hand sieves (2 mm mesh size) then stored in buckets to allow for individual identification until they were released back into the stream. Morphological identification of collected specimens was conducted by CB following Zuanon et al. (2015).

### Bioinformatic analysis

The bioinformatic analysis was completed using the *OBITools* metabarcoding package (Boyer et al., 2016), following the protocol described by Sales et al. (2021). *FastQC* was used to assess the quality of the reads and a length filter (command *obigrep*) was used to select fragments of 140-190 bp and to remove reads with ambiguous bases. SWARM was then used to compute sequence differences between aligned pairs of amplicons to delineate MOTUs (Mahé, Rognes, Quince, de Vargas & Dunthorn, 2014). The taxonomic assignment was conducted using the *ecotag* command, which works in two phases: initially, it performs a search of the assigned reference database to locate the sequence with the highest overall similarity to the query sequence; then the similarity value obtained from the first step is set as the threshold for searches of additional sequences, equal to or lower than that of the threshold value within the assigned database. Stringent filtering steps were applied to the final dataset to remove Molecular Operational Taxonomic Units (MOTUs)/reads originating from sequencing errors or contamination to avoid false positives for the library (Table S3). To reach this target, all non-fish reads were removed from the dataset, including non-target species (e.g., human and domestic species reads) and MOTUs that were likely to have been carried over from contamination. To remove putative contaminants, the maximum number of reads recorded in the controls (field collection, DNA extraction, and PCR blanks) were removed from all samples. Finally, all MOTUs with <10 reads were removed from the final dataset. The final taxonomic assignment was conducted according to current fixed general thresholds: MOTUs were assigned at species level when matching the reference sequence with >97% similarity as performed in previous studies in the Neotropics (Sales et al., 2020, 2021), at genus level with 95-97% similarity, at family level with 90-95% similarity, and the highest taxonomic level of order was attributed to MOTUs with less than 90% similarity matching the reference sequences.

### Data analyses

Given the difficulties of taxonomic assignment without complete reference databases, species identification was not possible from eDNA metabarcoding data and therefore MOTUs as opposed to species were used for all subsequent analyses. Replicates were pooled (nine water samples per site for the Ducke streams, two water samples for Aturiá and three for the Solimões river) before the following statistical analyses.

The MetacodeR package version 0.3.4 (Foster, Sharpton & Grünwald, 2017) was used to analyse the overall taxonomic diversity of the final eDNA dataset. A heat tree displaying the overall sample reads was produced to display the patterns of MOTU distribution of the eDNA data per taxonomic family. To visualise the magnitude of uncertainty of taxonomic assignments, a schematic phylogenetic tree was adapted from Betancur-R et al. (2017) representing the orders, families, genera and species detected by eDNA metabarcoding with the respective number of MOTUs assigned to each taxonomic group.

For the analysis of the diversity contained within the eDNA dataset (MOTU richness/α-diversity and β-diversity), the data were analysed with a presence/absence approach as suggested by Li et al. (2018). The α-diversity for the eDNA data (richness) was calculated as the total number of MOTUs found in each sample site and the β-diversity was obtained by the Jaccard dissimilarity index using the *vegdist* function in the vegan 2.5-2 package (Oksanen et al., 2013). Principal Coordinates Analysis (PCoA) was then used to investigate the relationship between distance and sites generated through the *cmdscale* function in the β-diversity matrix. The α-diversity analysis for the netting dataset was calculated as the total number of identified species found in each sample site and the β-diversity analysis was performed using the same method applied to the eDNA dataset. The results of the separate eDNA and netting β-diversity analyses were then superimposed to produce a final figure.

## RESULTS

### Evaluation of taxonomic resolution of the 12S rRNA fragment

Analysis of the taxonomic delimitation power of the 12S fragment targeted by the MiFish primers indicated a high rate of ambiguous identification (13.40% - 23.59%) for values ranging from 1% to 3%, with higher rates associated with increased threshold values. Optimum threshold analysis identified 0.1% up to 0.5% as intraspecific values with the lowest number of cumulative errors (Fig. 2A). Barcoding gap analysis revealed an extensive overlap between intraspecific and interspecific divergences, and only 133 species out of 264 were successfully recovered due to the absence of a barcoding gap (Fig. 2B). The overlap between intraspecific and interspecific genetic distances was particularly evident for species still lacking a full taxonomic description (e.g., *Trichomycterus* spp., *Hypostomus* spp. and *Harttia* spp.). Still, high divergence values (>3%) were also found for several other species including the Amazonian cichlids *Heros severus* (10.31%) and *Aequidens metae* (8.21%), and the loricariids *Hypostomus affinis* (12.3%), *H. plecostomus* (3.64%) and *Harttia carvalhoi* (3.57%).

**Figure 2.**
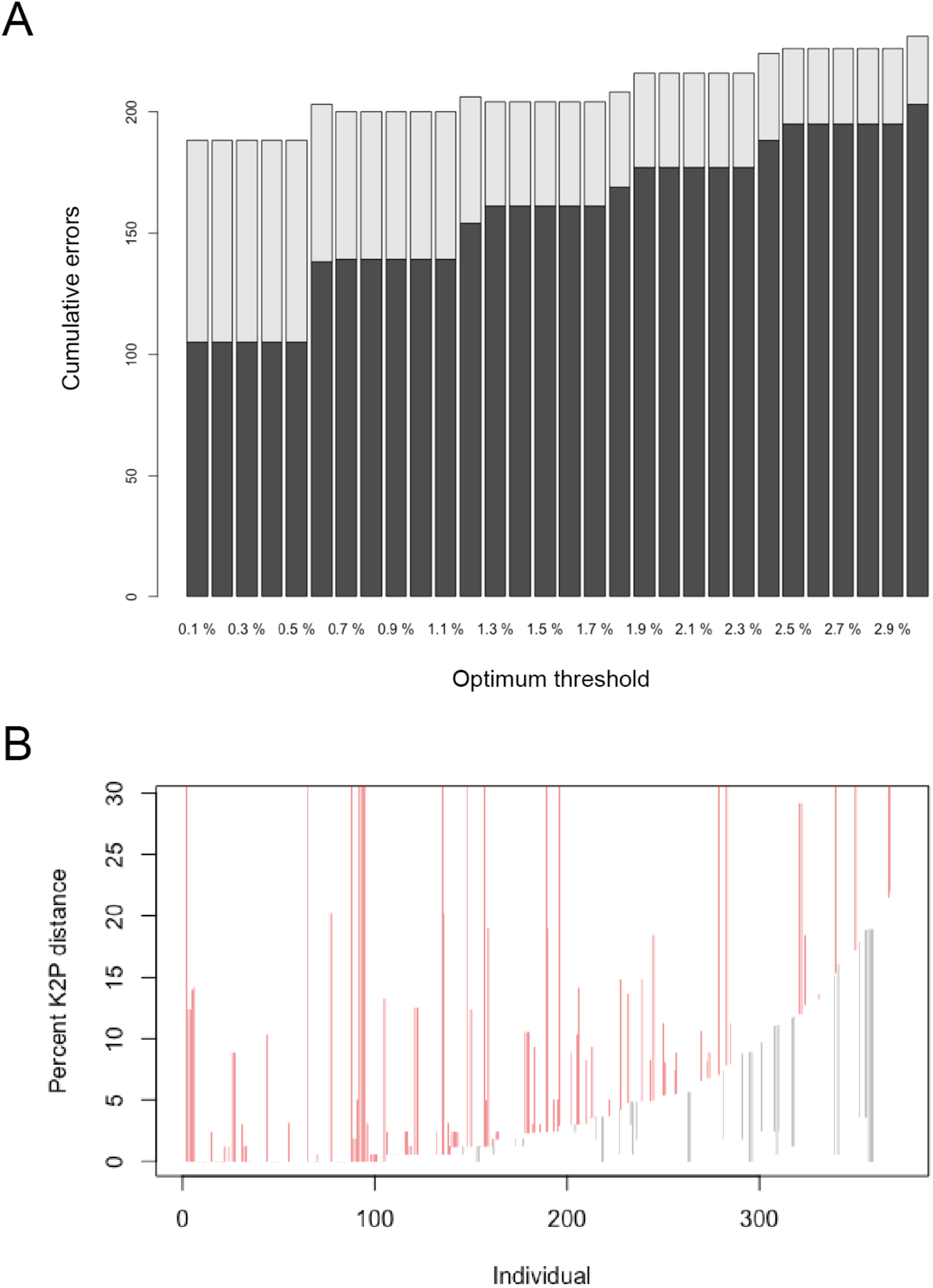
Optimum thresholds (A) and barcoding gap (B) analyses. Barplot (A) shows the false positive (light grey) and false negative (dark grey) rate of identification of species within the curated reference database at thresholds ranging from 0.1% to 3%. Line plot (B) of the barcode gap analyses for the 264 species (373 sequences) within the curated reference database. For each individual in the dataset, the grey lines represent the furthest intraspecific distance (bottom of line value), and the closest interspecific distance (top of line value) (i.e., presence of a barcode gap). The red lines show where this relationship is reversed (i.e., no barcode gap).

### eDNA data analysis

A total of 4,416,267 reads were obtained after trimming, merging, and length filtering during bioinformatic analysis (Table S5). Considering only MOTUs belonging to Actinopterygii, a final dataset containing 3,838,166 reads and 84 MOTUs was used for downstream analyses. A total of seven different taxonomic orders of fish were identified as a result of the taxonomic assignment: Characiformes, Cichliformes, Siluriformes, Gobiiformes, Synbranchiformes, Gymnotiformes, and Cypriniformes (Figs 3 and 4; Table S6). Taxonomic assignment was poor for the 84 recovered MOTUs; only four were identified to species level with the fixed general threshold, whereas 41 were assigned solely at the family level and 37 could only be attributed to the order level (Fig. 4). From the MOTUs identified to species level, one is known to occur in the Ducke Reserve (*Hoplias malabaricus*) and two have the genus present in this area (*Aequidens* and *Synbranchus*) and one has been detected only in the Aturiá (Site A; *Phreatobius* sp.). A MOTU was assigned to *Aequidens metae* (0.976 similarity) and *Aequidens pallidus* has been recorded from the Reserve. Another MOTU was assigned to *Synbranchus marmoratus* (0.976 similarity) but for the studied area, an as yet undescribed congeneric species (*Synbranchus* sp.) has been reported. A visualization of the fish community structure was obtained based on the taxonomic identification at the family level due to the difficulties of species identification. The assemblage structure depicted by the heat tree (Fig. 4) demonstrated a higher overall MOTU count for Characiformes (especially for the Anostomidae and Erythrinidae families) and Cichliformes (family Cichlidae). Furthermore, the heat trees evidenced the fish diversity recovered from eDNA samples due to the occurrence of 16 families among the 84 analysed MOTUs.

**Figure 3.**
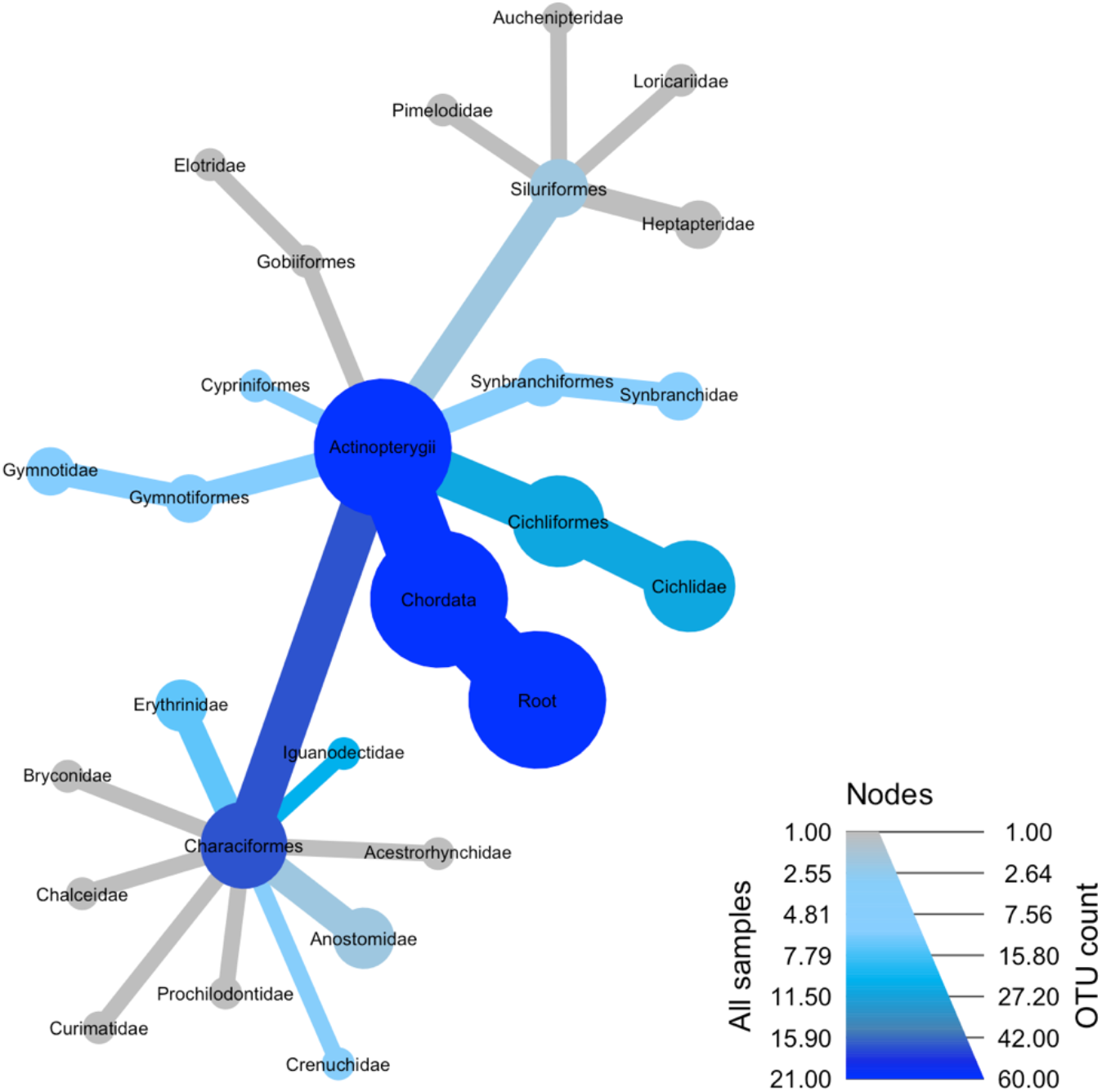
Heat tree displaying the fish diversity recovered for all sampling locations using eDNA metabarcoding. The blue colouration represents diversity identified from water samples, the darker the shade of blue, the more MOTUs detected for that taxonomic order.

**Figure 4.**
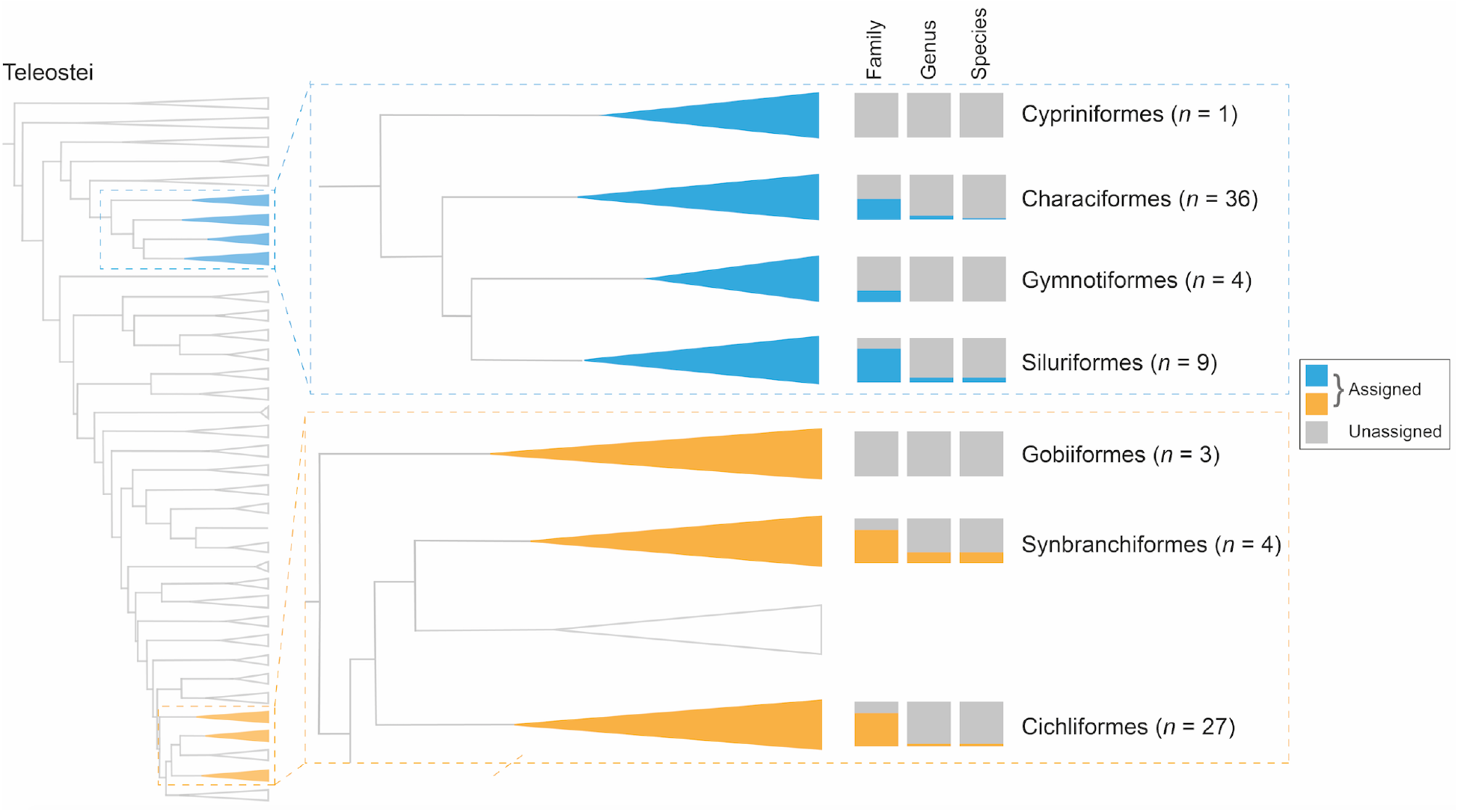
Represented orders within Teleostei (tree adapted from Betancur-R et al. 2017) to which the number of identified Molecular Operational Taxonomic Units (MOTUs) obtained from environmental DNA can be proportionally assigned to family, genus and species within each order.

### Alpha and beta diversity

The netting dataset revealed 18 different species in total (Figs 5A and 5B) with nine species for B1, 14 for B2, six for B3, and 11 for B4 (Figs 5B). Species richness for the eDNA data revealed a total of 32 unique MOTUs for the Ducke Reserve streams and a total of 46 MOTUs across the Ducke Reserve streams (B1: five MOTUs; B2: 19 MOTUs; B3: eight MOTUs and B4: 14 MOTUs; Fig. 6A). The eDNA dataset also revealed a total of 55 MOTUs for the Aturiá stream, 50 of which are unique to this sampling location with five of the MOTUs being shared with the other sampling locations. Three MOTUs (all of which are unique) were found in the Solimões (upper Amazon) river (Fig. S1).

**Figure 5.**
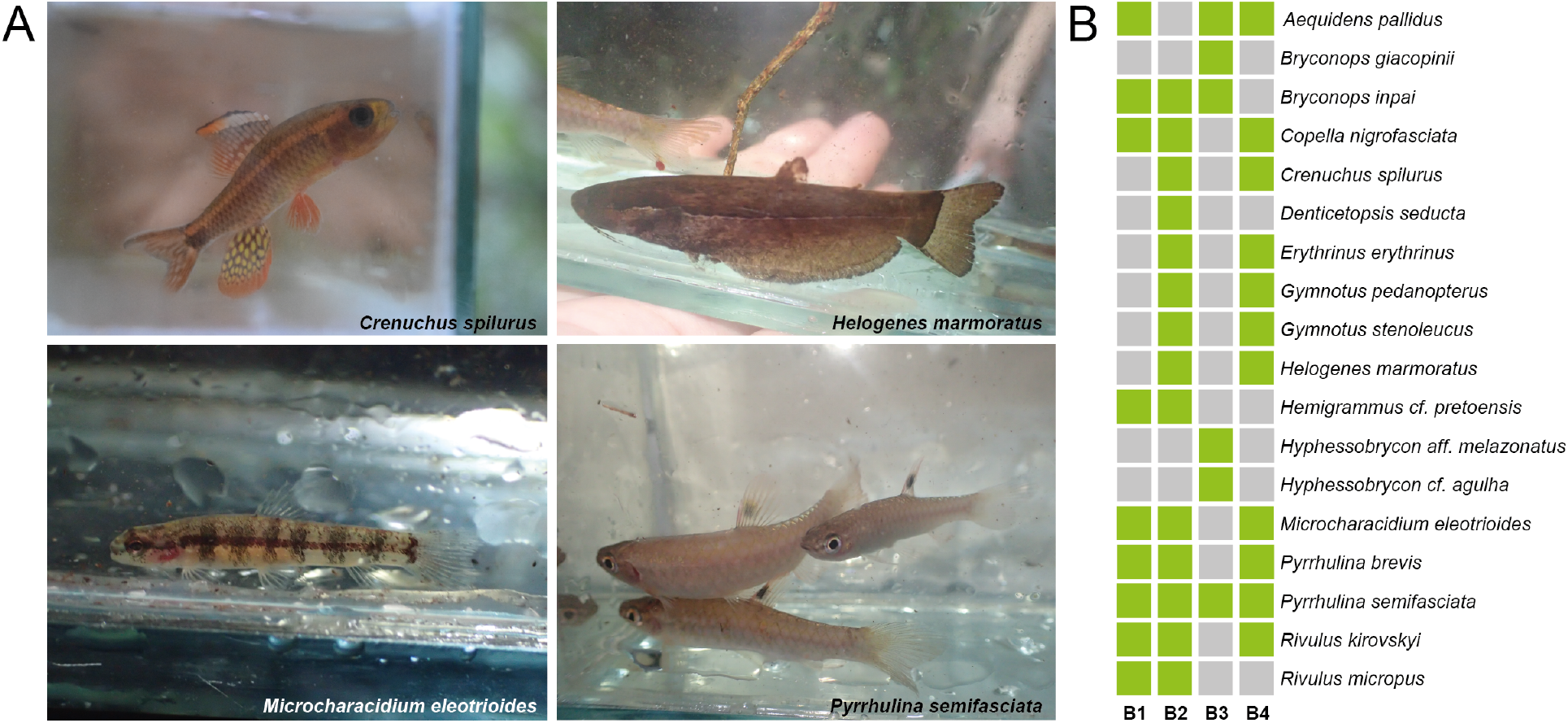
Examples of four species caught by netting in the Ducke Reserve (A), and presence (green) and absence (grey) of each of the 18 species caught in the four streams (B1-B4) within the Reserve (B).

**Figure 6.**
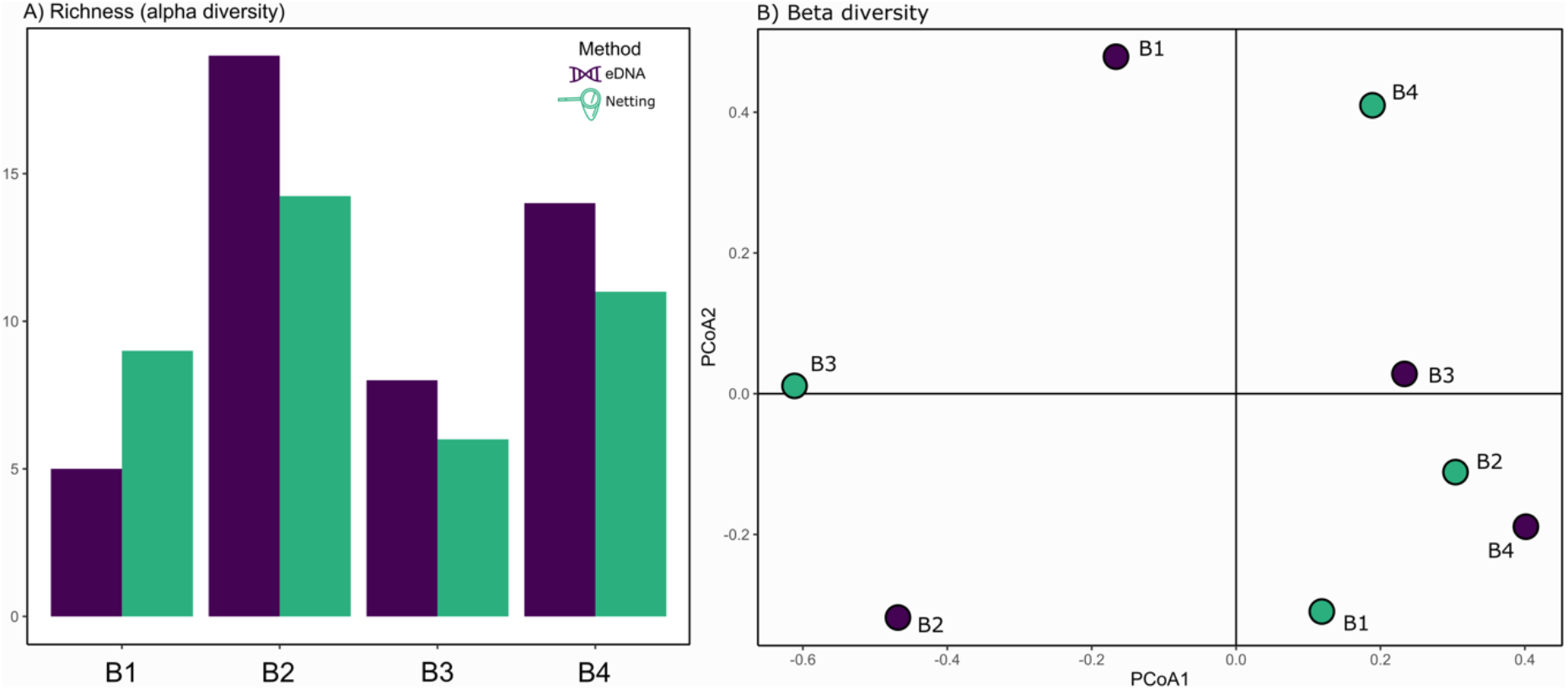
Species richness (A) and β-diversity inferred from separate PCoAs (B) based on eDNA MOTU data (purple) and netting data (green) from locations in the Ducke Reserve. See Fig. 1 for sampling locations.

β-diversity patterns inferred from Principal Coordinate Analysis (PCoA) showed a distinction between the Ducke Reserve streams and Aturiá and Solimões (Fig. S1). β-diversity also shows a clear distinction between the Aturiá stream and Solimões river. Within the Ducke Reserve, the four streams shared a total of 16 MOTUs and the β-diversity analysis shows a slightly more clustered pattern as opposed to the other two sampling locations in which Solimões shares no MOTUs with any other sampling location and the Aturiá shares only five MOTUs with the streams of the Ducke Reserve. Using only the sampling locations within the Ducke Reserve also demonstrated the distinction between the species compositions with considerable ordination distance between each point (Fig. 6B). Only sampling points B3 and B4 were slightly clustered, indicating that they share a more similar composition to each other compared to B1 and B2. Otherwise, large ordination distances separate all sampling locations within the Ducke Reserve. β-diversity analysis using the netting dataset again reveals marked distinctions between species compositions for all four analysed locations (Fig. 6B). However, in this case locations B1 and B2 showed the most similar compositions obtained through the netting data. β-diversity for the netting dataset did not show similar patterns compared with that obtained from the eDNA dataset (Fig. 6B).

## DISCUSSION

Understanding community compositions and species distributions within a target area is a crucial step to inform conservation management strategies. This requires cost-effective and robust methods for surveying biodiversity in a timely manner and eDNA metabarcoding has emerged as a promising method for surveying whole ecosystems (Lacoursière-Roussel et al., 2018; Murienne et al., 2019; Valdez-Moreno et al., 2019). As the use of genetic approaches to biodiversity monitoring relies on accurate, precise species identifications, the lack of appropriate taxonomic resolution and, consequently, detailed reference databases prevents the use of eDNA for biomonitoring from reaching its full potential. Herein, we demonstrate how the combination of these factors can significantly hinder the application of eDNA metabarcoding in monitoring fish communities in a megadiverse Neotropical system.

There is a knowledge gap regarding the optimum threshold for taxonomic assignments, as well as the taxonomic resolution/discriminative power of the gene fragments currently used in metabarcoding studies, particularly within megadiverse regions (Milan et al., 2020; Servis, Reid, Timmers, Stergioula & Naro-Maciel, 2020; Sales et al., 2021). Here, by performing a genetic-distance-threshold analysis, an optimum threshold value of 0.5% was obtained as a result of the lowest total cumulative errors (Fig. 2A). This value is in agreement with a previous study evaluating the taxonomic resolution of full length and mini-barcodes targeting the same gene (Milan et al., 2020, 0.55% for mini-barcodes), but contrasts with the standard threshold values (2-3%) currently widely applied for fish species delimitation based on various 12S rRNA fragments. Thresholds around 2% have been used in DNA barcoding studies targeting the mitochondrial COI region to delimit fish species and this has generally been extrapolated to eDNA surveys. However, using a fixed threshold (e.g., 2%) as a blanket value for studies in poorly described, species-rich areas can lead to incorrect conclusions regarding species richness. Consequences of this have been highlighted by Sales et al. (2018) where 306 fish DNA barcodes from the COI region showed that over one fifth had a mean intraspecific genetic divergence higher than 2%, flagging numerous possibilities of potential new MOTUs, cryptic species or errors originating from previous morphological identification of species.

For DNA barcodes to be considered effective and accurate, there should be a separation between intraspecific and interspecific divergences in the analysed marker, referred to as the “barcoding gap” (Meyer & Paulay, 2005). Analysis of the combined dataset investigating the presence of a barcoding gap revealed an extensive overlap between intraspecific variation and interspecific distances, leading to only 50% of species being recovered (Fig. 2B). The occurrence of no barcoding gap represents high levels of intra- and interspecific divergence within the analysed dataset. This suggests multiple possibilities: the presence of potential cryptic species (a common phenomenon for Neotropical fishes; Melo, Ochoa, Vari & Oliveira, 2016; Pugedo, de Andrade Neto, Pessali, Birindelli & Carvalho, 2016; Sales et al., 2018) errors within the reference database originating from poor taxonomy of previously identified species (Locatelli, McIntyre, Therkildsen & Baetscher, 2020); or poor taxonomic resolution of the marker used (Meyer & Paulay, 2005). Considering the high diversity of freshwater fish species in the Neotropics and consequently broad variation between intraspecific and interspecific genetic distances, caution should be taken when using universal cut-off values for species delimitation. As evidenced by the barcoding gap analysis, we stress that the application of a general threshold is not advisable, and values should be optimized considering the delimitation power of target fragments and the focal taxa. In order to achieve a robust taxonomic assignment, a thoughtful choice of primers is required. Although the 12S MiFish primers have undoubtedly been successfully applied in many studies and justifiably remain widely used (see Miya et al., 2020), it should be noted that it has been demonstrated that these primers might not provide an unambiguous identification of closely related species in some systems (Doble et al., 2020; Sales et al., 2021). Greater species level assignments can be achieved through the use of more specific primer sets (Doble et al., 2020), and/or by applying a multi-marker approach, taking into account that different markers/locus may show distinct taxonomic coverage and divergence ranges (Zhang et al., 2020).

Despite using a large dataset (373 sequences from 264 species), the results provided remain limited due to the lack of complete reference databases for Neotropical fish species. Not only are a low number of species are included but also, most of these are represented by just one sequenced specimen, hampering sound analyses regarding intraspecific genetic divergence and limiting further investigations. Another major impediment to the expansion of the reference data is the prevalence of efforts towards the construction of databases targeting very specific and short fragments, and more importantly, the lack of effort in making these sequences fully available in public depositories. For example, for Neotropical fish species, customised reference databases have been constructed targeting different fragments of the 12S gene (Cilleros et al. 2019; Milan et al., 2020; Sales et al., 2021) but in some cases only the fragment being analysed for eDNA has been made publicly available, even when a larger fragment of the gene has been sequenced for the reference database (Cilleros et al., 2019). This study highlights that the expansion of eDNA studies in megadiverse areas depends on better collaborative efforts focused on building more publicly available datasets.

The eDNA sampling successfully identified seven orders and 17 families of fish (Figs 3 and 4). If we consider the >99.5% threshold for species assignment based on the optimum threshold analyses presented above, only *Phreatobius* spp. would be assigned to species level. When considering a fixed general threshold of >97% for species assignment, only four species could be assigned (Fig. 4; Table S5). Two of these might represent different congeneric species (i.e., genera *Aequidens* and *Sybranchus*), possibly due to the lack of taxonomic resolution of the marker used (Yu et al., 2012). The potential for over and underestimations within our eDNA results is likely. We generated MOTUs using SWARM, aiming for an estimate of the overall biodiversity. While many MOTUs correspond to true biological species, some might be the result of PCR or sequencing errors, unidentified because of an incomplete reference database (Marques et al., 2019). The ambiguous identification values obtained (85-100%) may result in both the overestimation of true diversity (Reeder & Knight, 2009; Morgan et al., 2013) while bioinformatic clustering may have pooled closely related species, underestimating their number within certain taxonomic groups (Huse, Welch, Morrison & Sogin, 2010). This significantly hinders the information that we can draw from the eDNA dataset. Therefore, we used the information up to family level to depict the overall diversity detected in the eDNA data in which ecological inferences can be made. Despite underperforming eDNA accuracy regarding species identification, we could still draw important ecological information from the dataset. The high number of Characiformes MOTUs per site falls in line with previous studies in this region (Birindelli, Britski & Ramirez, 2020; Sales et al., 2021).

Species richness analysis of the eDNA and netting dataset within the Ducke Reserve reveals a similar pattern (Fig. 6A). Although sampling using eDNA detected slightly more MOTUs than species detected through the netting survey, it can be assumed that the 32 unique MOTUs obtained from the eDNA data might represent an overestimation of true diversity obtained (as outlined above). Species richness estimates obtained from netting surveys (Fig. 5) are reliable (but not infallible), given that each individual is assigned to a species with high confidence based on its morphological features (except for cryptic species). Nevertheless, while netting surveys produce accurate data, they cannot be applied to the required spatial scale in Neotropical basins. Furthermore, their success and reliability depend on the ability/expertise of the surveyor (labour intensive), the accessibility of the target area (challenging environments), and selectivity based on the deployed techniques’ ability to capture target species. Yet, when optimal conditions are met (complete reference database, and appropriate markers to name a few), eDNA will outperform traditional sampling, for its ability to detect species that are missed by the fishing gear deployed. In this study, the order Synbranchiformes, known to occur in the sampled area (Zuanon et al., 2015), was detected through eDNA, but not netting conducted at the same time. This species has been previously identified in the studied area and this highlights the randomness of net sampling, which could be due to the limited spatial and temporal scales covered by the netting survey, or, in this case, even by species behaviour. *Synbranchus* are nocturnal fish and can even survive buried up to three months, hence they can easily elude netting (Prestes-Carneiro & Béarez, 2017).

β-diversity analysis for both eDNA and traditional methods yielded contrasting patterns. While eDNA data showed similarities in compositions in the Ducke Reserve streams with slight clustering of sites B1 and B2, netting data suggest major differences between each sampling location (Fig. 6B). Barriers that hinder traditional surveying stemming from species size (affecting susceptibility of capture) and behavioural traits (i.e. shoaling/solitary, fast/slow, exposed/hidden) can be bypassed when analysing biodiversity on a genetic level (Evans, Shirey, Wieringa, Mahon & Lamberti, 2017). When utilising traditional methods, particularly in understudied areas, these barriers can create a bias towards species that are easier to capture, producing species-selective results (Holubová, Čech, Vašek & Peterka, 2019). A barrier that poses a great hindrance to biodiversity monitoring, and that is shared by both methods, is the presence of cryptic species (Beng & Corlett, 2020). Discerning between cryptic species may be difficult when species are morphologically similar and may result in the incorrect identification of a specimen. However, despite MOTUs being used as a proxy for species, and showing a positive correlation when used for taking ecological measures (e.g., β-diversity, Marques et al., 2019; Sales et al., 2021), results from the eDNA dataset should be used with caution as highlighted above. Therefore, prior to using these measures, eDNA metabarcoding and netting surveys merits further investigation in this Reserve.

Although eDNA metabarcoding potentially offers a means to assess biodiversity on a larger, more time-efficient scale, ensuring accuracy in results is critical. Evidently, eDNA metabarcoding as a biomonitoring tool in the Neotropical region is in its infancy, highlighted by the lack of appropriate reference databases. Herein, we argue that the commonly adopted threshold of >97% to assign MOTUs to species level is not optimal in megadiverse, understudied regions due to the likelihood of false positive or negative assignments. While this study shows that limited, albeit reliable, ecological inferences of biodiversity can be made with eDNA metabarcoding in this region (based on MOTU richness for example), it also highlights the need for significant investment in research aimed at improving availability of reference databases. We advocate for more transparency and collaboration within the research community, and recommend moving towards building whole mitochondrial genomes of specimens/species to identify multiple/other mini-barcodes and investigate order-specific taxonomic delimitation thresholds for future eDNA metabarcoding surveys of fishes and other vertebrates in the Neotropics (Milan et al., 2020; Sales et al., 2020, 2021).

## ACKNOWLEDGMENTS

The present study was carried out with all required permits (ICMBIO N. 54795-2, DEFRA 126191/385550/0). This project was partially funded by a University of Salford Internal Research Award awarded to CB, ADM, and IC. We are grateful to Rafael Estrela, Andrew Highlands and the undergraduate students of the University of Salford for assistance during fieldwork, Joseph Perkins for assistance during laboratory work and Sam Browett for help with bioinformatics. NGS also thanks FCT/MCTES for the financial support to CESAM (UID/AMB/50017/2019), through national funds. JSR and COC thank CNPq and the Biodiversity Research Consortium Brazil-Norway (BRC 16/19), for financial support through a PhD scholarship for COC and for consumable costs for sequencing. We also thank Guilherme Oliveira at the Instituto Tecnológico Vale, Belém, for laboratory access. The Ducke Reserve permanent stream plots are maintained by PELD-IAFA, PPBio-AmOC and INCT-CENBAM, all financed by the Ministry of Science, Technology and Innovation (MCTI) through CNPq and the Fundação de Amparo à Pesquisa do Estado de Amazonas (FAPEAM).

## AUTHOR CONTRIBUTIONS

NGS, ADM, CB and IC conceived, designed and acquired funding for the study. CB carried out the eDNA sampling and netting surveys. NGS performed the eDNA-based laboratory work and COC and JSR performed the species barcoding. NGS and JMJ carried out the bioinformatic analyses. JMJ, NGS, CB, IC and ADM analysed the data. JMJ, NGS and ADM wrote the manuscript, with all authors contributing to editing and discussions.

