## Supplementary Material for "eDNA in a bottleneck: obstacles to fish metabarcoding studies in megadiverse freshwater systems"

### TABLE OF CONTENTS


| | |
|---|---|
| **Table S1.** List of fish species sequenced including the samples identification number (ID) and the species record number in the Taxonomy database (NCBI). | Page 3 |
| **Table S2.** Orders included in the combined enhanced dataset with the total number of families and species analysed. | Page 5 |
| **Table S3.** Each sampling location within the Ducke reserve and respective coordinates. | Page 5 |
| **Table S4.** List of analysed samples, including sample code, location, distance and side of sampling, date of collection, date of DNA extraction and final DNA concentration. | Page 6 |
| **Table S5.** Filtering steps removing all MOTUs/reads originating from sequencing errors or contamination and respective number of reads retrieved at each stage. | Page 9 |
| **Table S6.** Taxonomic orders of fish detected through eDNA metabarcoding with the total number of MOTUs detected at each taxonomic level. | Page 9 |
| **Figure S1.** MOTU richness (A) and beta-diversity inferred from a PCoA (B) in all six sites sampled for environmental DNA. See Fig. 1 for sampling locations. | Page 10 |

### APPENDIX

#### Study site

The Reserve is a designated 100 $km^2$ area protecting continuous and non-isolated rainforest established by the National Institute of Amazon Research (INPA) in the 1960s and is located north-east of Manaus (Somavilla & Oliveira, 2017). At the time of establishment, Manaus had a population of around 173,000 people in an area of approximately 3,000 hectares (Silva & Silva, 1993). Today, the estimated population size has risen to >2,000,000 inhabitants spread over an increased area of >11,000,000 hectares (Silva & Galvão, 2017). The accelerated population growth of Manaus (over 40 years) has resulted in encroachment into the reserve (Peres & Terborgh, 1995), increasing the risk of forest fragmentation, pollution of water systems, and deforestation.

**Table S1.** List of fish species sequenced including the samples identification number (ID) and the species record number in the Taxonomy database (NCBI).

| ID | SPECIES | Taxonomy ID NCBI |
|---|---|---|
| O8B83 | *Acestrorhyncus falcatus* | 639289 |
| 11B07 | *Aequidens pallidus* | 2478851 |
| 11AF4 | *Ammocryptocharax elegans* | 1569731 |
| 10A03 | *Ancistrus* aff. *hoplogenys* | NO |
| 11A74 | *Aphyocharacidium* sp. | NO |
| 10A47 | *Apistogramma agassizi* | 284746 |
| 11A39 | *Apistogramma hippolytae* | NO |
| 10C41 | *Brycon melanopterus* | 686976 |
| 11A20 | *Bryconops* cf. *caudomaculatus* | 1463024 |
| 11A38 | *Bryconops giacopinii* | NO |
| 11C52 | *Bryconops inpai* | NO |
| 11A86 | *Carnegiella strigata* | 642546 |
| 09C40 | *Characidium* cf. *pteroides* | NO |
| 11A69 | *Characidium pellucidum* | NO |
| 11B05 | *Copella nigrofasciata* | NO |
| 09F37 | *Crenicichla inpa* | NO |
| 11A40 | *Crenicichla alta* | 238234 |
| 10A49 | *Crenuchus spilurus* | 909854 |
| 11AC03 | *Denticetopsis epa* | NO |
| 1559 | *Eigenmania* aff. *macrops* | 749476 |
| 10A07 | *Erythrinus erythrinus* | 754154 |
| 11A70 | *Farlowella* sp. | NO |
| 11A67 | *Gnathocharax steindachneri* | 42533 |
| 10C97 | *Gymnorhamphychthys rondoni* | 679697 |
| 10A08 | *Gymnotus carapo* | 94172 |
| 08C51 | *Helogenes marmoratus* | 337700 |
| 11A89 | *Hemigrammus* cf. *pretoensis* | NO |
| 11A71 | *Hemigrammus vordwinkleri* | NO |
| 09B19 | *Hoplias malabaricus* | 27720 |
| 11A29 | *Hyphessobrycon* cf. *agulha* | NO |

| | | |
|---|---|---|
| 10B82 | *Hypopygus lepturus* | 36692 |
| 11A59 | *Iguanodectes geisleri* | 767257 |
| 09D13 | *Ituglanis amazonicus* | 2023032 |
| 10B85 | *Mastiglanis asopos* | NO |
| 09C08 | *Microcharacidium weizmani* | NO |
| 09C82 | *Microsternarchus bilineatus* | 36688 |
| 10A56 | *Monocirrhus polyacanthus* | 302772 |
| 11A33 | *Myoglanis koepckei* | NO |
| 11B06 | *Nannostomus marginatus* | NO |
| 11A51 | *Odontocharacidium aphanes* | NO |
| 11A95 | *Parotocinclus* sp. | NO |
| 10B53 | *Phenacogaster pectinatus* | NO |
| 11B86R | *Pyrrhulina* sp. *(brevis/semifasciata group)* | 42609 |
| 11A79 | *Rineloricaria* sp. | 52092 |
| 11A07 | *Satanoperca* sp. | NO |
| 10A29 | *Sternopygus macrurus* | 77841 |
| 10B16 | *Synbranchus* sp. | NO |

* The "NO" in the Taxonomy ID NCBI column indicates species that are currently not included in the NCBI taxonomy database and therefore do not now hold a registration ID number.

**Table S2.** Orders included in the combined enhanced dataset with the total number of families and species analysed.

| Taxonomic order | Family count | Species count |
|---|---|---|
| Characiformes | 19 | 122 |
| Cyprinodontiformes | 1 | 5 |
| Gymnotiformes | 4 | 7 |
| Osteoglossiformes | 1 | 1 |
| Perciformes/Cichliformes | 3 | 38 |
| Pleuronectiformes | 1 | 1 |
| Siluriformes | 5 | 89 |
| Synbranchiformes | 1 | 1 |

**Table S3.** Each sampling location within the Ducke reserve and respective coordinates.

| Sampling locations | Latitude | Longitude |
|---|---|---|
| Acará _L2 (B1) | -2.9361 | -59.9628 |
| Acará _L3 (B2) | -2.9447 | -59.9550 |
| Acará _L4 (B3) | -2.9508 | -59.9569 |
| Barro Branco (B4) | -2.9264 | -59.9703 |
| Solimões (C) | -3.1289 | -59.8964 |
| Aturiá (A) | -2.0358 | -61.1494 |

**Table S4.** List of analysed samples, including sample code, location, distance and side of sampling, date of collection, date of DNA extraction and final DNA concentration.

| Sample | Location | Distance | Side | Date Collection | Date Extraction | DNA [ng/uL] |
|---|---|---|---|---|---|---|
| **DK01** | BLANK_Acara_L2 (B1) | NA | NA | 18.01.19 | 17.04.19 | 0.2 |
| **DK02** | BLANK_Acara_L4 (B3) | NA | NA | 17.01.19 | 17.04.19 | 0.4 |
| **DK03** | Acara_L4 (B3) | 0 | Right | 17.01.19 | 17.04.19 | 1.6 |
| **DK04** | Acara_L4 (B3) | 0 | Left | 17.01.19 | 17.04.19 | 1.4 |
| **DK05** | Acara_L4 (B3) | 0 | Centre | 17.01.19 | 17.04.19 | -0.2 |
| **DK06** | Acara_L4 (B3) | 25 | Right | 17.01.19 | 17.04.19 | 1.5 |
| **DK07** | Acara_L4 (B3) | 25 | Left | 17.01.19 | 17.04.19 | 0.8 |
| **DK08** | Acara_L4 (B3) | 25 | Centre | 17.01.19 | 17.04.19 | 2.5 |
| **DK09** | Acara_L4 (B3) | 50 | Right | 17.01.19 | 17.04.19 | 0.8 |
| **DK10** | Acara_L4 (B3) | 50 | Left | 17.01.19 | 17.04.19 | 1.2 |
| **DK11** | Acara_L4 (B3) | 50 | Centre | 17.01.19 | 17.04.19 | 0.7 |
| **DK12** | Acara_L2 (B1) | 0 | Right | 18.01.19 | 17.04.19 | 8 |
| **DK13** | Acara_L2 (B1) | 0 | Left | 18.01.19 | 17.04.19 | 4.4 |
| **DK14** | Acara_L2 (B1) | 0 | Centre | 18.01.19 | 17.04.19 | 2.4 |
| **DK15** | Acara_L2 (B1) | 25 | Right | 18.01.19 | 17.04.19 | 5.1 |
| **DK16** | Acara_L2 (B1) | 25 | Left | 18.01.19 | 17.04.19 | 0.2 |
| **DK17** | Acara_L2 (B1) | 25 | Centre | 18.01.19 | 17.04.19 | 3 |
| **DK18** | Acara_L2 (B1) | 50 | Right | 18.01.19 | 17.04.19 | 5 |
| **DK19** | Acara_L2 (B1) | 50 | Left | 18.01.19 | 17.04.19 | 3.4 |
| **DK20** | Acara_L2 (B1) | 50 | Centre | 18.01.19 | 17.04.19 | 1.8 |
| **DK21** | Barro Branco (B4) | 0 | Left | 15.01.19 | 17.04.19 | 1.9 |
| **DK22** | BLANK_extraction | NA | NA | NA | 17.04.19 | -0.4 |

| | | | | | | |
|---|---|---|---|---|---|---|
| **DK23** | BLANK_Acara_L3 (B2) | NA | NA | 16.01.19 | 18.04.19 | 1.3 |
| **DK24** | BLANK_Barro Branco (B4) | NA | NA | 15.01.19 | 18.04.19 | 1.9 |
| **DK25** | Barro Branco (B4) | 0 | Right | 15.01.19 | 18.04.19 | 3.6 |
| **DK26** | Barro Branco (B4) | 0 | Centre | 15.01.19 | 18.04.19 | 4.1 |
| **DK27** | Aturiá (A) | NA | NA | 21.01.19 | 18.04.19 | 21.3 |
| **DK28** | Aturiá (A) | NA | NA | 21.01.19 | 18.04.19 | 9.6 |

| Sample | Location | Distance | Side | Date Collection | Date DNA Extraction | DNA [ng/uL] |
|---|---|---|---|---|---|---|
| **DK29** | Barro Branco (B4) | 25 | Right | 15.01.19 | 18.04.19 | 8 |
| **DK30** | Barro Branco (B4) | 25 | Left | 15.01.19 | 18.04.19 | 5.3 |
| **DK31** | Barro Branco (B4) | 25 | Centre | 15.01.19 | 18.04.19 | 1.9 |
| **DK32** | Barro Branco (B4) | 50 | Right | 15.01.19 | 18.04.19 | 1.6 |
| **DK33** | Barro Branco (B4) | 50 | Left | 15.01.19 | 18.04.19 | 6.6 |
| **DK34** | Barro Branco (B4) | 50 | Centre | 15.01.19 | 18.04.19 | 3.7 |
| **DK35** | Acara_L3 (B2) | 0 | Right | 16.01.19 | 18.04.19 | 1 |
| **DK36** | Acara_L3 (B2) | 0 | Left | 16.01.19 | 18.04.19 | 3.3 |
| **DK37** | Acara_L3 (B2) | 0 | Centre | 16.01.19 | 18.04.19 | 0.9 |
| **DK38** | Acara_L3 (B2) | 25 | Right | 16.01.19 | 18.04.19 | 3.5 |
| **DK39** | Acara_L3 (B2) | 25 | Left | 16.01.19 | 18.04.19 | 4 |
| **DK40** | Acara_L3 (B2) | 25 | Centre | 16.01.19 | 18.04.19 | 5.1 |
| **DK41** | Acara_L3 (B2) | 50 | Right | 16.01.19 | 18.04.19 | 2.8 |
| **DK42** | Acara_L3 (B2) | 50 | Left | 16.01.19 | 18.04.19 | 6.2 |
| **DK43** | Acara_L3 (B2) | 50 | Centre | 16.01.19 | 18.04.19 | 2 |
| **DK44** | Solimoes_1 (C) | NA | NA | NA | 18.04.19 | 5.8 |

| | | | | | | |
|---|---|---|---|---|---|---|
| **DK45** | Solimoes_2 (C) | NA | NA | NA | 18.04.19 | 2.6 |
| **DK46** | Solimoes_3 (C) | NA | NA | NA | 18.04.19 | 6.2 |
| **DK47** | BLANK_extraction | NA | NA | NA | 18.04.19 | 0.2 |

**Table S5.** Filtering steps removing all MOTUs/reads originating from sequencing errors or contamination and respective number of reads retrieved at each stage.

| Filtering Steps | Total |
|---|---|
| **Total Reads** | **4,416,267** |
| After removing reads from the blanks | 4,393,789 |
| After removing all non-fish reads | 3,839,422 |
| After removing all total reads <10 | 3,838,166 |

**Table S6.** Taxonomic orders of fish detected through eDNA metabarcoding with the total number of MOTUs detected at each taxonomic level.

| **ORDER** | **Number of MOTUS** | | | |
|---|---|---|---|---|
| | **ORDER** | **FAMILY** | **GENUS** | **SPECIES** |
| Characiformes | 36 | 16 | 3 | 1 |
| Cichliformes | 27 | 20 | 1 | 1 |
| Cypriniformes | 1 | 0 | 0 | 0 |
| Gobiiformes | 3 | 0 | 0 | 0 |
| Gymnotiformes | 4 | 1 | 0 | 0 |
| Siluriformes | 9 | 7 | 1 | 1 |
| Synbranchiformes | 4 | 3 | 1 | 1 |
| **Total** | 84 | 47 | 6 | 4 |

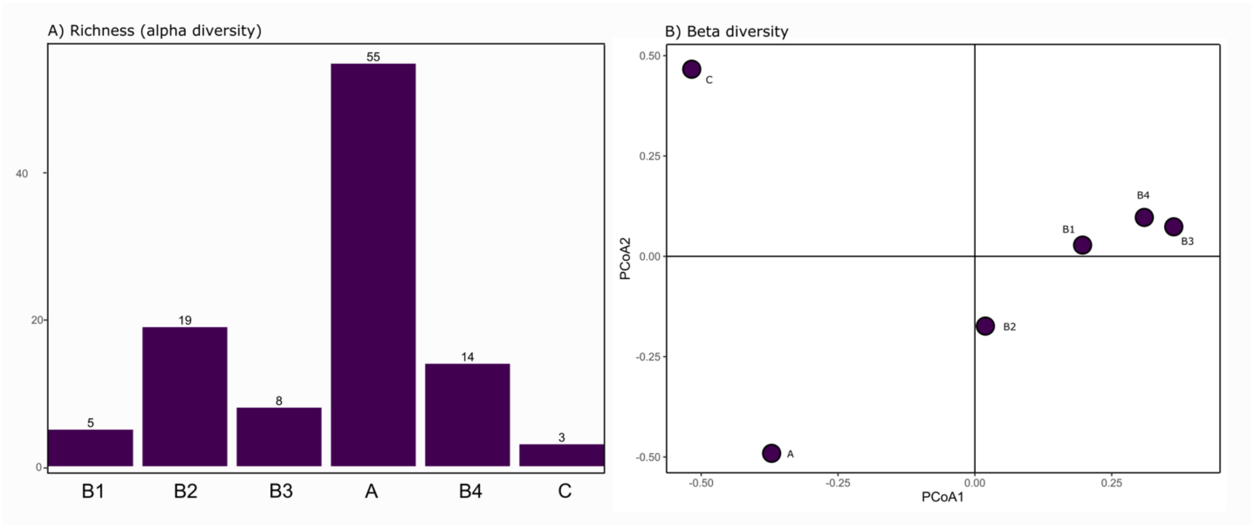


**Figure S1.** MOTU richness (A) and beta-diversity inferred from a PCoA (B) in all six sites sampled for environmental DNA. See Fig. 1 for sampling locations.
